# MTAP loss correlates with an immunosuppressive profile in GBM and its substrate MTA stimulates alternative macrophage polarization

**DOI:** 10.1101/329664

**Authors:** Landon J. Hansen, Rui Yang, Karolina Woroniecka, Lee Chen, Hai Yan, Yiping He

## Abstract

Glioblastoma (GBM) is a lethal brain cancer known for its potent immunosuppressive effects. Loss of *Methylthioadenosine Phosphorylase* (*MTAP*) expression, via gene deletion or epigenetic silencing, is one of the most common alterations in GBM. Here, we show that MTAP loss in GBM cells is correlated with differential expression of immune regulatory genes. In silico analysis of gene expression profiles in GBM samples revealed that low *MTAP* expression is correlated with reduced proportions of γδT cells, fewer activated CD4 cells, and an increased proportion of M2 macrophages. Using in vitro macrophage models, we found that methylthioadenosine (MTA), the metabolite that accumulates as a result of MTAP loss in GBM cells, promotes the immunosuppressive alternative activation (M2) of macrophages. We show that this effect of MTA on macrophages is independent of IL4/IL3 signaling, is mediated by the adenosine A_2B_ receptor, and can be pharmacologically reversed. This study suggests that MTAP loss in GBM cells contributes to the immunosuppressive microenvironment, and that *MTAP* status should be a factor for consideration in understanding GBM immune states and devising immunotherapy-based approaches for treating *MTAP*-null GBM.

## INTRODUCTION

Immunotherapy possesses enormous potential for treating cancer and has reshaped the way we understand and treat certain cancer types (1,2). Despite recent progress, however, the promise of immunotherapy-based approaches for treating brain tumors, in particular high grade glioblastoma (GBM), remains to be fully realized (3-5). GBM is the most common and lethal brain tumor, with a dismal median survival of 12-15 months from the time of diagnosis (6). It is also a cancer characterized by its immune-suppressive nature. It has been well-established that GBM cells actively employ multiple strategies to escape immune surveillance and to create an immunosuppressive microenvironment (7-9). As such, to fully harness the power of immunotherapy for GBM requires better understanding and more effective strategies for countering the tumors’ immunosuppressive effects.

Recent genomic studies have provided insights into the molecular mechanisms of GBM pathogenesis, revealing the most commonly mutated genes in tumor cells (10,11). Gliomas can be classified based on their genetic alterations and gene expression profiles into subtypes that predict tumor characteristics and patient prognosis (12). Further studies, in both glioma and other types of cancer, have associated genetic alterations with tumor cells’ evasion of immune surveillance and manipulation of the immune microenvironment, providing new rationale for tailoring treatment based on cancer cells’ genetic composition (13-15). As an example, recent discoveries have linked *IDH1* mutations in glioma to immune evasion through interference with immune activation pathways, providing new opportunities for devising highly specific immunological treatments (16-19). Thus, identifying additional cancer-specific mutations that confer a similar immune escape/ suppression-based advantage to glioma cells will likely lead to new rationale for immunotherapeutic designs.

One of the most common genetic alterations in GBM, occurring in approximately 50% of all cases, is the homozygous deletion or epigenetic silencing of methylthioadenosine phosphorylase (*MTAP*) (10,20). We have recently demonstrated that *MTAP* deletion is associated with increased tumorigenesis and with shortened disease-free survival in GBM patients (Hansen, et al, submitted). MTAP is a metabolic enzyme that functions in the salvage pathway of adenine and methionine, and loss of MTAP results in the accumulation of its direct metabolite substrate, methylthioadenosine (MTA), in both intracellular and environmental compartments (21-23). MTA is known to be functionally active within cells as an inhibitor of methyltransferases (22,23). This metabolite has also been shown to suppress cell proliferation via different, cellular context-dependent mechanisms, including targeting the Akt signal pathway and interfering with intracellular protein methylation in T cells (24), or acting through adenosine receptors on the cell surface of melanoma cell lines (25). Studies on mechanism of pathogen-induced host inflammatory responses have linked MTA to downregulation of TNFα production by macrophages through engaging adenosine receptors (26), and have revealed a role of MTA in controlling host inflammation response, such that MTA has been used as an immunosuppressive drug for treating colitis, liver inflammation, brain inflammation and autoimmunity in animal models (27-29). Lastly, MTA has been shown to be elevated in the blood of septic patients (30) and in the urine of children with severe combined immunodeficiency (SCID) (31), further supporting its role in regulating the immune response.

In this study, we investigated the link between MTAP loss in GBM cells and the GBM microenvironment. We show that *MTAP* expression correlates with the expression of genes regulating innate or adaptive immune response in both experimental cell models and in GBM samples. We reveal that in GBM tissues, low expression level of MTAP is associated with altered immune cell populations indicative of a more immunosuppressive context. In particular, we provide evidence that MTAP loss-induced MTA accumulation stimulates M2 alternative macrophage activation. We illustrate that this effect of MTA on macrophages is independent of IL4/IL3 signaling, is mediated by the adenosine A_2B_ receptor and STAT3 signaling, and is distinct from the actions of adenosine. Finally, we show that the MTA-induced alternative macrophage activation can be pharmacologically reversed. These results provide a basis for blocking adenosine A_2B_ receptor signaling as a strategy to potentiate immunotherapy targeting *MTAP*-null GBMs.

## RESULTS

### MTAP loss in GBM cells is associated with an immunosuppressive gene expression profile

We established patient-derived GBM cell lines, verified *MTAP* status, and used the affymetrix gene expression microarray to characterize their global gene expression profile (L. Hansen, et al., submitted). Analysis of differentially regulated genes (unpaired ANOVA comparing *MTAP* WT vs *MTAP* null cell lines) revealed a list of downregulated inflammatory pathways, including the KEGG pathway of antigen processing and presentation (**Fig. 1A**). One group of genes common to all these pathways are the human leukocyte antigen (HLA) genes, which were found to be consistently downregulated in tumor cells that lack MTAP (**Fig 1B**). We compiled a gene set with all HLA genes included in the gene expression microarray (Supporting data Table 2) and performed gene set enrichment analysis (GSEA) as previously described (32) to examine the correlation of this set of genes with *MTAP* status. GSEA confirmed that *MTAP* null cells displayed markedly lower expression of HLA genes, including HLA Class I and both α and β chains of HLA class II (**Fig. 1C**). Consistent with our finding of differentially expressed inflammatory pathways, GSEA using gene sets of inflammatory cytokines showed that *MTAP* null cells had reduced expression of inflammatory cytokines compared to *MTAP* WT cells (**Fig. 1D**). Examination of individual genes showed that while several regulators of the adaptive immune response were affected (*IL2, CTLA4*, and *CD44*), most differentially expressed genes were associated with the innate immune response, including those that regulate monocyte/macrophage proliferation, differentiation and activation, such as *CSF1, IL13, IL34, IL37, ALOX5*, and toll-like receptor related genes, *TLR4, TIRAP*, and *LY96* (**Fig. 1E**). Collectively, these results illustrate that *MTAP* expression correlates with numerous immune-regulatory genes in a manner suggesting that loss or reduction of *MTAP* expression contributes to the immune-suppressive nature of GBM cells.

**Figure 1.**
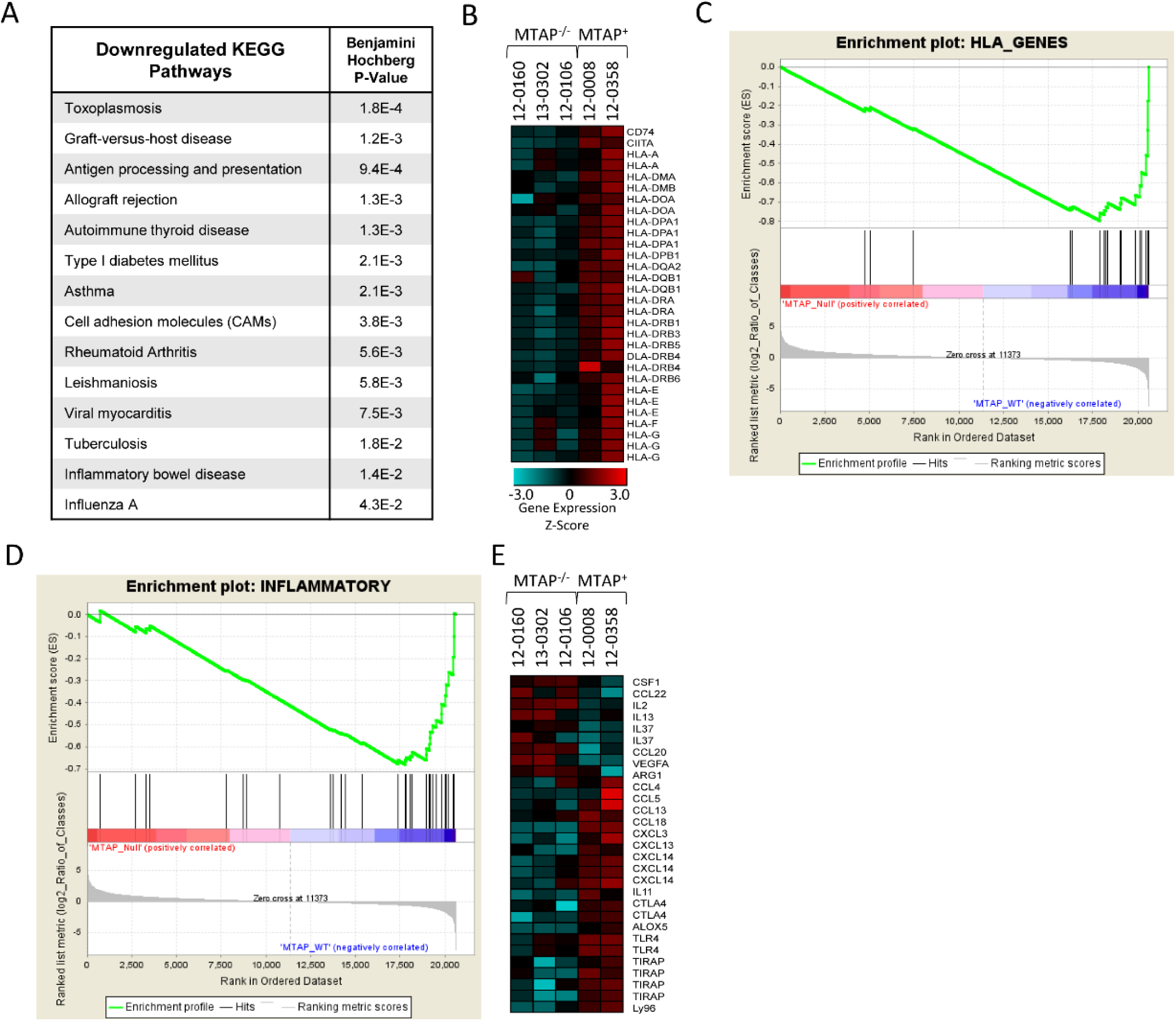
MTAP loss is correlated with downregulated inflammatory genes in GBM cells. **(A)** DAVID pathway analysis of 3,000 most downregulated genes in *MTAP* null cells compared to *MTAP* expressing cells reveals downregulated pathways related to inflammatory processes. **(B)** Heatmap showing expression of HLA genes is downregulated in *MTAP* null patient-derived GBM cell lines compared to *MTAP* WT cell lines. All transcripts with t test *P* value <0.05 comparing the *MTAP* deleted and *MTAP* WT cell lines are included. 17/25 HLA transcripts are significantly downregulated (29 unique probes). No HLA transcripts are significantly upregulated in *MTAP* deleted cells. **(C)** Gene set enrichment analysis using a list of all HLA genes in the Affymetrix 2.0 plus microarray demonstrates significant enrichment of HLA genes in *MTAP* WT compared to *MTAP* deleted cell lines, Normalized Enrichment Score = −2.18, FDR q-value < 0.001. **(D)** Pathway analysis using a list of inflammatory cytokines (see supporting data for gene set) demonstrates enrichment of inflammatory gene expression in *MTAP* WT compared to *MTAP* deleted cell lines, Normalized Enrichment Score = −2.12, FDR q-value < 0.001. **(E)** Heatmap comparing expression of individual transcripts between the *MTAP* deleted and *MTAP* WT cell lines.

### Loss of MTAP expression is associated with an immunosuppressive molecular profile in GBM

To evaluate the relevance of the above *in vitro* findings to GBM, we took advantage of The Cancer Genome Atlas (TCGA) large gene expression dataset (33,34) to test the link between *MTAP* expression and the immune status of GBM tumor samples. First, we analyzed expression of individual inflammatory mediators/ regulators from the TCGA GBM dataset. Consistent with the findings from the *in vitro* models, this analysis showed that low *MTAP* expression was correlated with lower expression of cytokines known to promote inflammatory responses (e.g. *CCL4, CCL5, IL2, TNF, IFNG*), and with higher expression of genes known to be involved in regulating monocyte/macrophage activation, including *TGFB1, CHI3L1, CHII3L2, HRH1, TREM2*, and *P2RY13* (Supporting Fig. S1).

To test the association of *MTAP* expression with the immune cell profile in GBM we employed CIBERSORT, a recently developed method for deconvoluting gene expression data and estimating immune cell fractions in human cancers (35-37). This method was chosen for several reasons: (i) it has been successfully used for analyzing a large number of samples across multiple cancer types (36), (ii) it is readily compatible with the microarray-based platform of the largest available GBM expression dataset, and (iii) this approach’s unique consideration of data normalization and noise control can be particularly helpful when analyzing GBM samples which are known for their heterogeneity and diffusiveness (38). To validate the application of this approach to glioma samples, we first evaluated the TCGA low-grade glioma dataset (39) by testing the immune cell components in tumors with wild-type or mutant *IDH1/2*, since the effect of *IDH* mutations on glioma immune suppression has been well defined (17-19). In particular, it has been shown that *IDH* mutation is correlated with lower levels of CD8+ and CD4+ T cells and lower macrophage infiltration (17,18). CIBERSORT analysis of the low grade glioma gene expression dataset from TCGA (n=530 samples) (39) revealed a profile consistent with these previously reported observations (Supporting Fig. S1). Interestingly, CIBERSORT also revealed a previously unknown effect of *IDH* mutation on numbers of macrophages, mast cells, plasma cells, and B cells, with a higher representation of each cell type, except macrophages, in *IDH*-mutant gliomas (Supporting Fig. S2).

We extended the same analysis to the GBM dataset from TCGA (385 GBM patient samples) (10) to examine the link between MTAP loss and the GBM immune microenvironment. As MTAP alteration can occur through homozygous gene deletion or epigenetic silencing, we compared the two groups of patients with the highest and lowest *MTAP* expression (upper and lower quartiles, n=96 each) (Supporting Fig. S3). This analysis revealed reduced fractions of activated CD4 T cells in tumors with low *MTAP* expression (**Fig. 2A**). Additionally, two types of innate immune cells were identified as differentially represented between these two groups of tumors. First, there was a lower fraction of γδT cells in the low-*MTAP* expressing tumors (**Fig. 2B**); and second, a significantly higher fraction of M2 macrophages were observed in low-*MTAP* tumors compared to high MTAP-expressing tumors (**Fig. 2C**).

**Figure 2.**
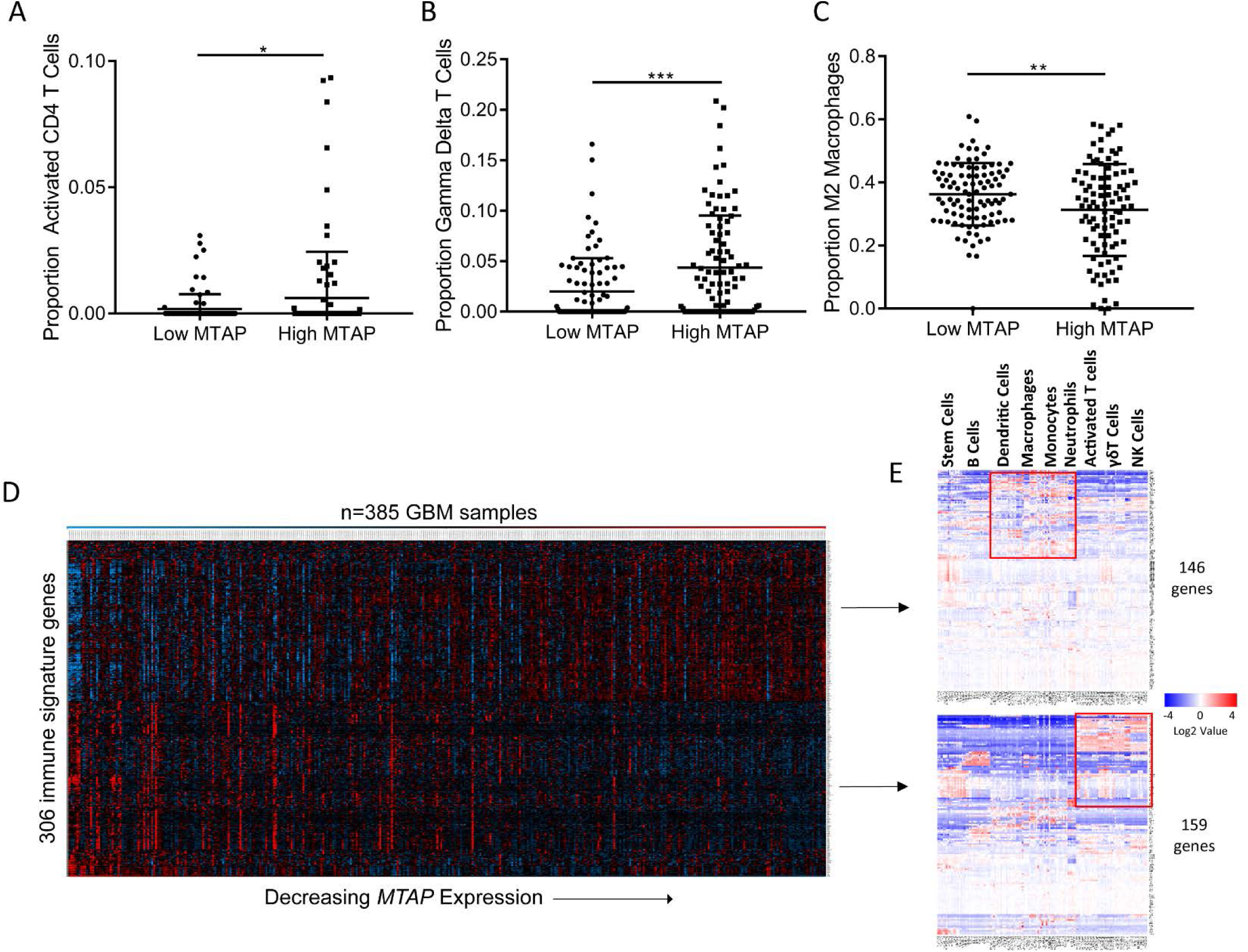
MTAP loss is correlated with an immunosuppressive gene signature in GBM samples. **(A)** Comparison of upper and lower *MTAP* expression quartiles (n=96 each category) using CIBERSORT analysis of GBM gene expression microarray data shows lower activated CD4 T cells in tumors with low *MTAP* expression. **(B)** GBM samples with the lowest quartile *MTAP* expression have fewer gamma delta T cells. **(C)** GBM samples with low *MTAP* expression have higher average proportions of M2 macrophages, n=96 each category. **(D)** Heatmap showing expression of 306 immune signature genes that are differentially expressed between the lower and upper *MTAP* expression quartiles in GBM patients. The gene list was derived from 782 immune-specific genes identified by Charoentong et al. Only 641 of the 782 genes were contained in the GBM microarray data set. Of the 641 genes in the GBM dataset, 306 genes were differentially expressed based on MTAP status, and were included in the heatmap. **(E)** Clustered gene sets from (D) were analyzed using data from the immunologic genome project (https://www.immgen.org), which uses a compendium of microarray data (366 microarrays/37 studies) from specific immune cell types to display which immune cells express the genes of interest. Clusters representing different immune cell types are marked at the top. Genes expressed in the low MTAP samples are more representative of macrophages, monocytes, dendritic cells, while genes expressed in high MTAP samples are more representative of activated CD4 cells, gamma delta (γδ) T cells, and natural killer (NK) cells. t test *P* value * P < 0.05, ** P < 0.005, *** P < 0.0005.

To corroborate the findings from CIBERSORT analysis, we utilized a recently developed platform for identifying individual immune cell types based on a dataset of 366 microarrays compiled from multiple independent studies (40). This data set was previously used to generate a list of 782 genes that are specific to immune cell subpopulations (i.e. not expressed in tumor cells or normal tissue) (40). We analyzed this list of genes in the TCGA GBM dataset and found that among the 641 of these genes for which expression data was available, 306 of them were differentially expressed between the MTAP-low and MTAP-high GBM populations (upper quartile vs lower quartile, n=96 each group, t test *P* value < 0.05), with an even distribution between the two groups (146 genes upregulated in MTAP-low tumors, 159 genes upregulated in MTAP-high tumors) (**Fig. 2D**). When these two derivative groups of genes were linked to their respective immune cell subtypes using the previously defined immune cell gene signatures (40), we found that the genes upregulated in MTAP-low tumors were predominantly associated with macrophages and monocytes, while those genes upregulated in MTAP-high tumors were expressed in activated CD4 T cells, natural killer cells, and γδ T cells (**Fig. 2E**; Supporting Figs. S4 and S5). Collectively, the findings from our various analyses on the gene expression profiles of low MTAP and high MTAP expressing GBM samples suggest a reduction in immune-reactive T cells (41) and an increase in immunosuppressive M2 macrophages (42) in samples with low *MTAP* expression, supporting the notion that MTAP deficiency is linked to a more immunosuppressive GBM microenvironment.

### MTA stimulates alternative activation of macrophages

Among the aforementioned immune cell types that were found to be differentially represented between *MTAP*-low versus *MTAP*-high GBMs, the potential impact of *MTAP* loss on macrophage populations was particularly intriguing. Macrophages are the most abundant immune cell type in GBM, representing as many as half of all cells in the tumor mass (43), where they are known to play an immunosuppressive/protumoral role (42). Macrophages can be regulated by adenosine signaling (44), which potentiates the effect of cytokines in promoting alternative macrophage activation through adenosine A_2_ receptors (45). Studies focusing on pathogenic mechanism and treatment of pathogen-induced host inflammatory responses have found that MTA can suppress Lipopolysaccharide (LPS)-induced expression of inflammatory response genes, such as *TNF*, via adenosine A_2_ receptors (26,46). These indications, together with our finding of a higher fraction of M2 macrophages in low-*MTAP*-expressing GBMs as described above, led us to hypothesize that MTA potentiates M2 macrophage activation through adenosine receptor signaling.

We tested this hypothesis using the well-established murine RAW 264.7 macrophage cell line model (45,47). As expected, treatment of RAW 264.7 cells with M2 macrophage-inducing Th2-type cytokines, interleukin (IL)-4 and IL-13, drove the cells toward the alternative type of activation, as demonstrated by the upregulated expression of *Arginase 1* (*Arg1*), a canonical marker for M2 macrophages (48) (**Fig. 3A**). When MTA was used in combination with IL-4/IL-13, induction of *Arg1* expression was dramatically augmented (**Fig. 3B**). Remarkably, when MTA was used as a single agent to treat the cells, upregulated expression of *Arg1* was also observed (**Fig. 3C**). This effect was not limited to RAW 264.7 cells as it was also observed in the murine BV-2 microglial cell line and in the human THP-1 cell line (**Fig. 3D**, Supporting Fig. S6A). Notably, the effect of MTA on macrophages was distinct from that of IL-4 or IL-13, as treatment of cells with MTA attenuated the IL-4/IL-13-induced expression *Pparg* and *Mrc1*, marker genes associated with the M2a macrophage subtype (**Fig. 3E**; Supporting Fig. S6B) (49,50). Also distinct from the effect of IL-4 or IL-13 was that MTA activated the expression of *Vegfa*, a classic feature of the M2d subtype of alternatively activated macrophages (**Fig. 3F** and **G**) (50). In agreement with this finding from the *in vitro* model, we revisited the GBM gene expression data (TCGA) and validated that in GBM patients *VEGFA* expression was significantly higher in samples with low MTAP expression (**Fig. 3H**). Collectively, these results suggest that MTA promotes alternative macrophage activation resembling the M2d subtype and that this process likely occurs in MTAP deficient GBMs.

**Figure 3.**
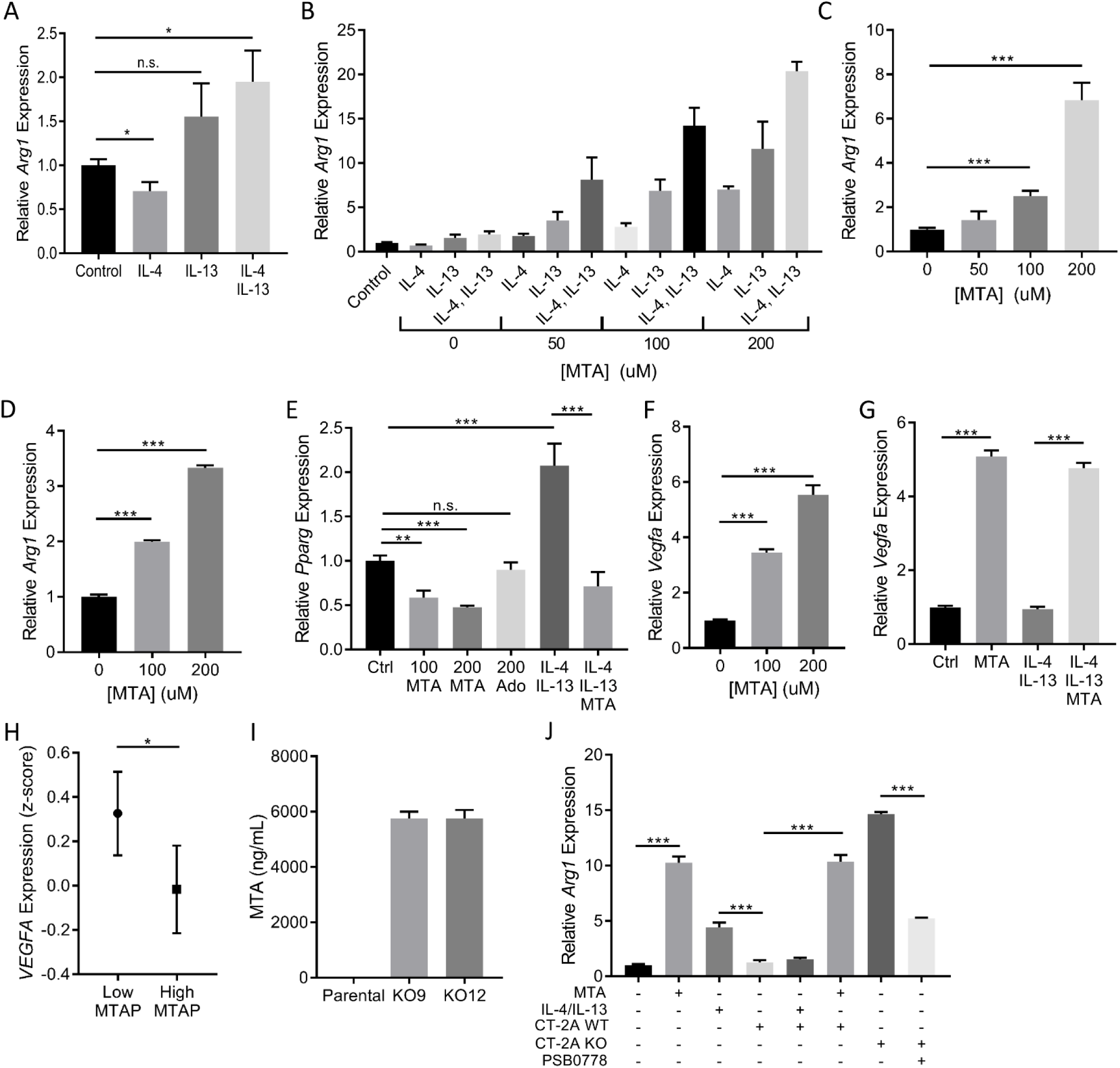
MTA promotes upregulation of M2 macrophage marker genes. **(A)** RAW 264.7 cells were treated with IL-4 and IL-13 individually or in combination (5ng/mL for each cytokine) for 12 hours before total RNA was prepared for *Arg1* expression analysis (all gene expression was measured by RT-qPCR. **(B)** RAW 264.7 cells were treated with the indicated cytokines as in (A), together with different doses of MTA, after which relative *Arg1* expression was determined as above. **(C)** RAW 264.7 or **(D)** BV-2 cells were treated with different doses of MTA for 12 hours and *Arg1* expression was measured. **(E-G)** RAW 264.7 cells were treated with the indicated cytokines with or without MTA for 12 hours, and total RNA was isolated for measuring expression of (E) *Pparg* and (F, G) *Vegfa*. Ado=adenosine. **(H)** TCGA human GBM microarray data were used for analyzing *VEGFA* expression in tumors with low or high *MTAP* expression, n=96 patients per group. (**I**) MTA was measured in spent cell culture media from CT-2A parental and MTAP knockout cell lines using liquid chromatography tandem mass spectrometry (LC-MS/MS). (**J**) Raw 264.7 cells were exposed to media collected from parental or *MTAP* knockout CT-2A cells for 12 hours before *Arg1* gene expression was measured by RT-qPCR. All experiments were repeated independently. All statistical comparisons were performed using an unpaired student’s t-test; * = *P*<0.05, ** = *P*<0.005, *** = *P*< 5×10^−4^, n.s. = not significant.

To validate that the phenotype induced by exogenously administered MTA is relevant to the GBM extracellular microenvironment, we generated an isogenic *MTAP*-null derivative of the murine GBM cell line CT-2A. As expected, CRISPR-mediated homozygous deletion of *MTAP* in CT-2A cells led to the accumulation of MTA in the culture media (**Fig. 3I**). We then tested the response of macrophages to spent media from the *MTAP*-null cell line as a way of simulating the tumor microenvironment. Exposure of RAW 264.7 cells to spent media from *MTAP*-null CT-2A cells indeed induced *Arg1* expression. While exposure to the spent media from parental CT-2A (*MTAP* wildtype) cells had a minimal effect on *Arg1* expression, the addition of exogenous MTA to the conditioned media was able to stimulate an alternative activation response (**Fig. 3J**). Together, these results support the hypothesis that MTA in the extracellular compartment of *MTAP*-null GBM cells can work in concert with known M2 macrophage-stimulating cytokines in altering tumor associated macrophages, and/or directly promoting the alternative activation of macrophages as a single agent.

### Regulation of macrophage activation by MTA requires the adenosine A2B receptor and STAT3

Previous work has shown that the action of adenosine in modulating the innate immune response to cytokine signaling is achieved through the adenosine A_2_ receptors (50). We tested whether the effects of MTA are also mediated through these receptors using specific antagonists of A_2A_ and A_2B_ receptors. We found that an A_2A_ receptor antagonist, Istradefylline, had only minimal effect on MTA-induced *Arg1* expression. In contrast, an antagonist of the A_2B_ receptor, PSB0778, while failing to block the effect of IL-4/IL13 as expected, potently attenuated the expression of *Arg1* induced by MTA (**Fig. 4A**), an effect which was also seen in the BV-2 cell model (**Fig. 4B**). Furthermore, PSB0778 also blocked the induction of *Arg1* expression by the spent media of *MTAP*-null GBM cells (Fig. 3J), and attenuated other MTA-stimulated M2 marker genes, including *Vegfa, Timp1*, and *IL10*, while countering the inhibitory effect of MTA on *Mrc1* expression (**Fig. 4C**; Supporting Fig. S6B). These results suggest that MTA acts through the adenosine A_2B_ receptor to regulate macrophage activation.

**Figure 4.**
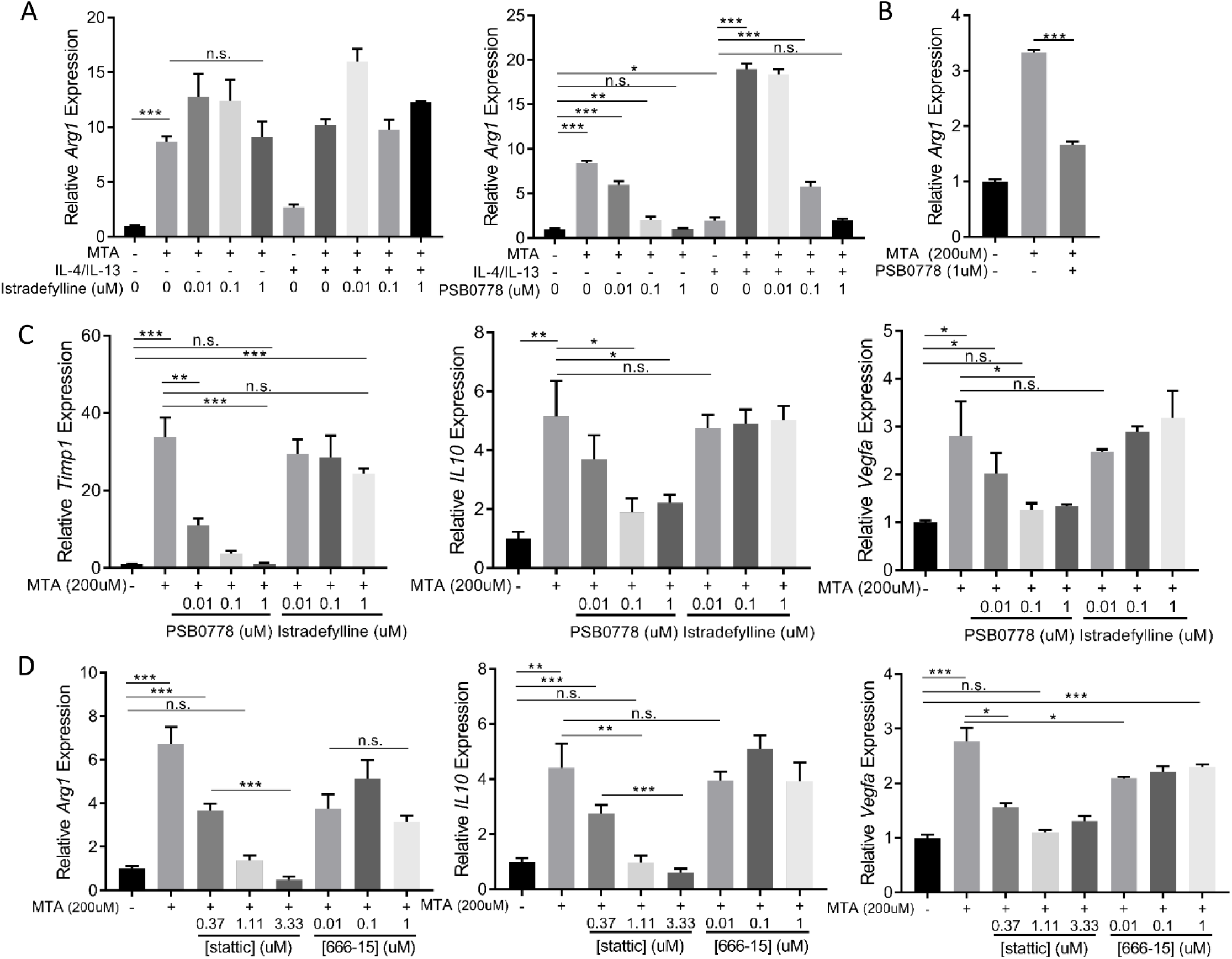
The impact of MTA on M2 macrophage marker genes is blocked by inhibition of the A2B adenosine receptor and Stat3. **(A)** RAW 264.7 cells were treated with MTA and/or IL-4/IL-13 for 12 hours and simultaneously with (left) A_2A_ receptor antagonist Istradefylline or (right) A_2B_ receptor antagonist PSB0778 and expression of *Arg1* was determined by RT-qPCR. **(B)** The BV-2 microglial cell line was treated with MTA with or without A_2B_ receptor antagonist PSB0778 for 12 hours and *Arg1* expression was measured by RT-qPCR. **(C)** RAW 264.7 cells were treated with MTA, with or without A_2B_ receptor antagonist PSB0778 or A_2A_ receptor antagonist Istradefylline for 12 hours and expression of *Timp1, IL10* and *Vegfa* were determined by RT-qPCR. **(D)** RAW 264.7 cells were treated with MTA, with or without STAT3 inhibitor Stattic or CREB inhibitor 666-15 for 12 hours, and expression of *Timp1, IL10*, and *Vegfa* were determined by RT-qPCR. All statistical comparisons were done using an unpaired student’s t-test; * = *P*<0.05, ** = *P*<0.005, *** = *P*< 5×10^−4^, n.s. = not significant.

The essential role of the adenosine A_2B_ receptor in mediating the effect of MTA led us to ask whether a sustained contribution/function of this signaling pathway is required for maintaining the resultant macrophage activation state. To address this question, we treated the cells with the A_2B_ receptor antagonist at a delayed time point following treatment with M2-inducing cytokines and/or with MTA. We found that treatment of the already activated macrophages with the adenosine A_2B_ receptor antagonist was still able to abolish *Arg1* expression (Supporting Fig. S6C and D), suggesting the effect of MTA can be pharmacologically reversed.

To further illuminate the downstream mediators of MTA-stimulated A_2B_ receptor signaling, we tested the role of transcriptional regulators CREB and STAT3, as these are reported to function downstream of adenosine receptors and are known to regulate *Arg1, Vegfa*, and *IL10*, among other genes (51-55). We found that the inhibition of STAT3 completely abrogated the MTA-induced expression of all three genes. In contrast, inhibition of CREB had only marginal or no effect on MTA-induced gene expression (**Fig. 4D**). These results suggest that STAT3 is a necessary component of M2d macrophage polarization induced by MTA-stimulated A_2B_ receptor signaling.

### MTA and adenosine activate distinct yet overlapping signaling pathways

As both adenosine and MTA signal through adenosine receptors, we sought to determine if there were any quantitative or qualitative differences in their effect on macrophage activation. We treated RAW 264.7 macrophages with equimolar concentrations of adenosine or MTA and tested the expression of macrophage activation markers in response to each metabolite. To our surprise, we found that MTA much more potently stimulated expression of *Arg1, Vegfa*, and *Timp1* than did adenosine (**Fig 5A**). In addition, MTA was unique in its ability to suppress *Pparg* expression (**Fig. 5B**). Furthermore, while MTA and adenosine similarly upregulated *IL10* expression (**Fig. 5C**), the impact of adenosine on *IL10* (and *Timp1*) was impervious to PSB0778, suggesting adenosine was acting through a different receptor than MTA to induce expression of these genes (**Fig. 5A** and **5C**). Notably, the impact of MTA on *IL10* was also only partially reduced by A_2B_ receptor antagonism (Fig 4C and 5C).

**Figure 5.**
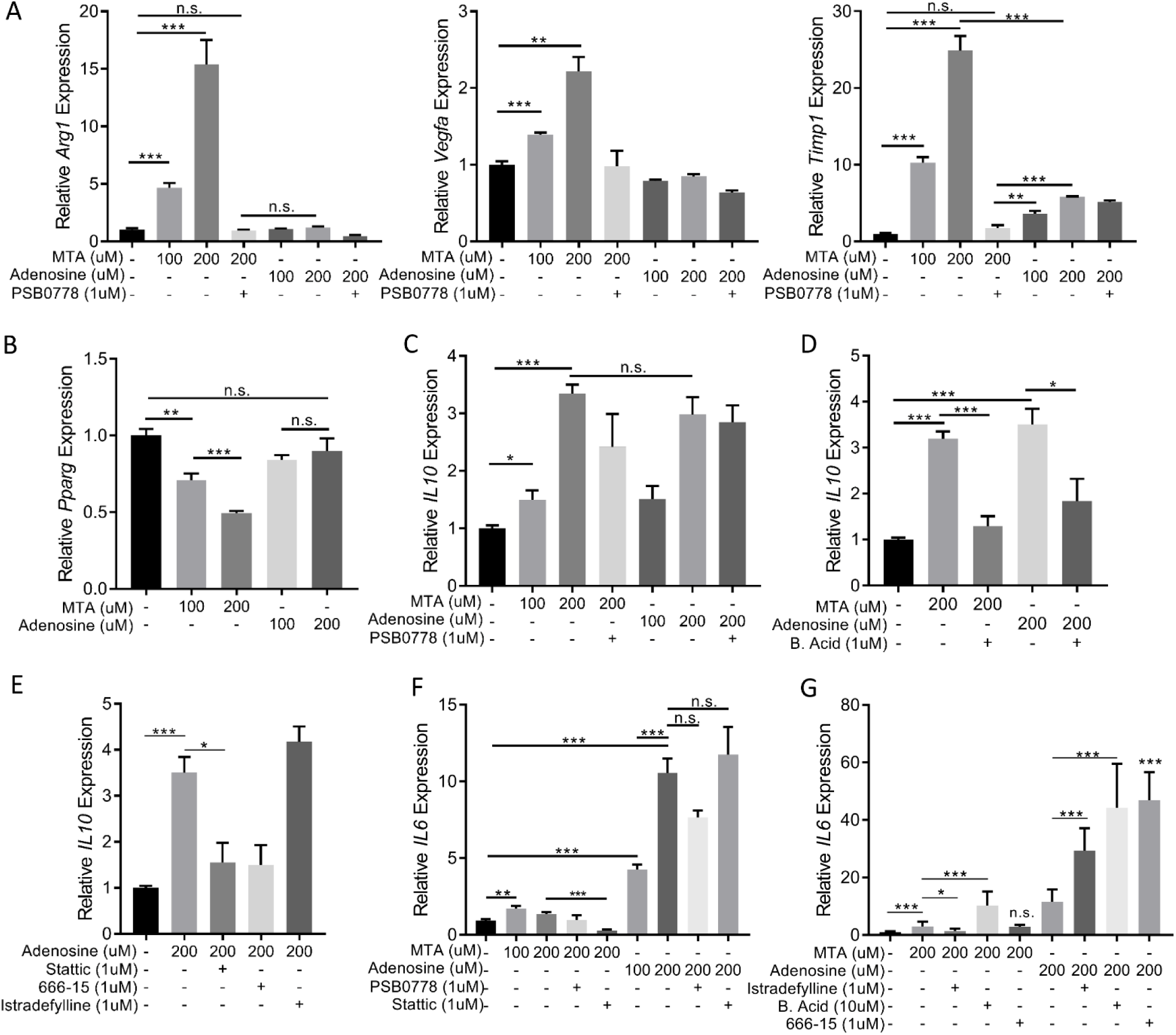
MTA and adenosine have distinct effects. **(A)** RAW 264.7 cells were treated with the MTA or adenosine with or without A_2B_ receptor antagonist PSB0778 for 12 hours and *Arg1, Vegfa, Timp1* expression were measured by RT-qPCR **(B)** RAW 264.7 cells were treated with MTA or Adenosine for 12 hours and *Pparg* expression was measured by RT-qPCR. **(C and D)** Raw 264.7 cells were treated with MTA or adenosine for 12 hours with or without (C) A_2B_ receptor antagonist PSB0778 or (D) C/EBP inhibitor betulinic acid (B. Acid) as indicated and *IL10* expression was measured by RT-qPCR. **(E)** *IL10* expression was measured by RT qPCR following adenosine treatment with or without STAT3 inhibition (Stattic), CREB inhibition (666-15), or A_2A_R antagonist Istradefylline. **(F and G)** RAW 264.7 cells were treated with MTA or adenosine for 12 hours with or without the indicated inhibitors and *IL6* expression was measured by RT-qPCR. All samples were collected 12 hours after treatment. All statistical comparisons were performed using an unpaired student’s t-test; * = *P*<0.05, ** = *P*<0.005, *** = *P*< 5×10^−4^, n.s. = not significant.

We then tested the role of downstream mediators in mediating the response of MTA and adenosine signaling. We found the effect of MTA and adenosine on *IL10* expression were both effectively blocked by betulinic acid, an inhibitor of transcription factor C/EBP (CEBPA/CEBPB) (**Fig. 5D**), consistent with reports of C/EBP being a critical regulator of *IL10* expression (56) and indicating a partial convergence of the adenosine and MTA signaling pathways through this transcription factor. Furthermore, similar to what we had observed with MTA (Fig. 4D), induction of *IL10* expression by adenosine was also blocked by stattic, indicating adenosine-stimulated *IL10* requires STAT3 (**Fig. 5E**). However, whereas MTA-induced expression of *IL10* was independent of CREB (Fig. 4D), addition of CREB inhibitor 666-15 effectively blocked *IL10* expression in response to adenosine (**Fig. 5E**), indicating that in the downstream signaling pathways there are distinct mechanisms responsible for MTA and adenosine-mediated responses in macrophages.

To further illuminate the distinct/overlapping roles of MTA and adenosine, we investigated the expression of *IL6*, another important regulator of macrophage activation (57,58) that has been reported to be upregulated in response to adenosine signaling (59-63). We found that adenosine much more potently upregulated *IL6* than did MTA, and that while inhibition of STAT3 decreased *IL6* expression in the context of MTA treatment, STAT3 inhibition had no effect after adenosine treatment (**Fig. 5F**). Furthermore, adenosine-mediated upregulation of *IL6* expression was potentiated by A_2A_ receptor antagonist istradefylline, as well as by inhibition of CREB or C/EBP, while the effect of MTA was only potentiated by the C/EBP inhibitor, betulinic acid, suggesting there are different negative feedback signaling pathways for MTA and adenosine (**Fig. 5G**). Thus, while the effect of MTA and adenosine on *IL6* expression again showed some overlap (both metabolites upregulated *IL6*, though adenosine to a greater extent, and the addition of C/EBP inhibition further increased *IL6* expression in response to both), the dramatic differences in *IL6* regulation in response to MTA or adenosine highlight the quantitative distinction in the degree to which different receptors and pathways are activated by these two metabolites.

The finding that adenosine upregulates both *IL10* and *IL6* expression while MTA stimulates only *IL10* provides a potential explanation for the distinct gene expression patterns resulting from MTA or adenosine treatment (Fig. 5A and B). It is recognized that IL-10 has a more anti-inflammatory effect than IL-6, despite both cytokine receptors activating Jak1/STAT3 (54,64,65). This difference in inflammatory signaling is thought to be due to feedback regulation of IL-6 signaling by STAT3-induced Socs3 (65), whose expression was upregulated following both MTA and adenosine treatment in our model (Supporting Fig. S6E). IL-10 has been shown to be essential for the anti-inflammatory response, through its opposition of IL-6 and its role in suppressing inflammatory cytokines (66), thus raising the possibility that activation of *IL10* in response to MTA may potentiate the anti-inflammatory effect of adenosine receptor signaling.

Collectively these results suggest that MTA and adenosine each engage a distinct balance of adenosine receptor signaling, resulting in a divergent pathway activation and gene expression profiles (**Fig. 6)**. This unique effect of MTA on macrophage M2 activation is not shared by adenosine, requires the adenosine A_2B_ receptor, STAT3, and C/EBP, and indicates an immunosuppressive effect of MTA accumulation in MTAP deficient GBM.

**Figure 6.**
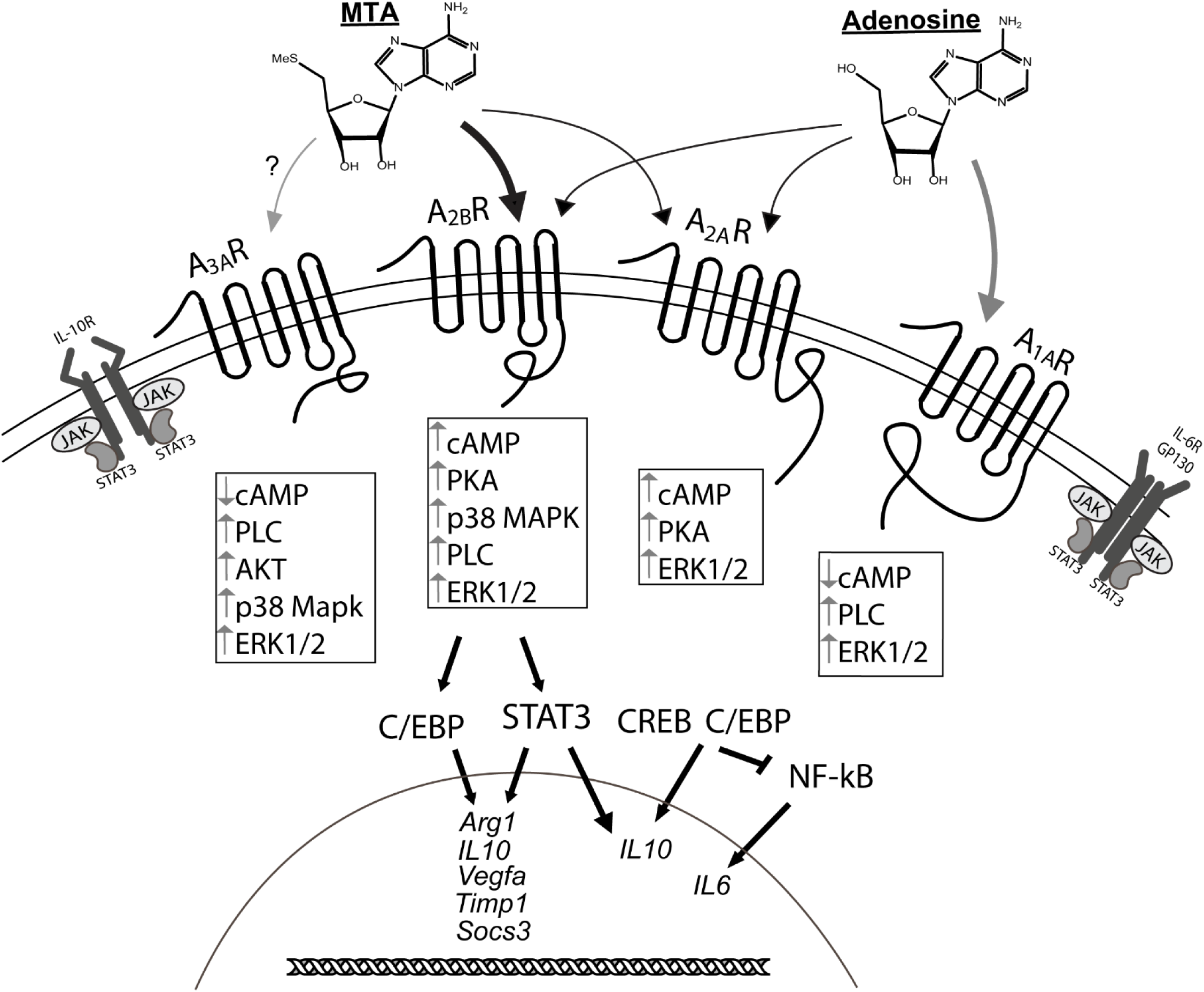
Simplified proposed mechanism of MTA and adenosine-mediated signaling in macrophages. Using small molecule inhibitors specific to the A_2B_ receptor (PSB0778) and the A_2A_ receptor (Istradefylline) as well as inhibitors of STAT3 (Stattic), CREB (666-15) and C/EBP (Betulinic Acid), we measured expression of M2 macrophage-associated marker genes following treatment with MTA or adenosine. The ability of MTA to upregulate M2 macrophage marker genes was dependent on A_2B_ receptor signaling, and required C/EBP and STAT3. Upregulation of *IL10* expression by adenosine was blocked by inhibition of STAT3, C/EBP or CREB, while inhibition of the A_2A_ receptor, C/EBP, or CREB resulted in upregulation of *IL6* expression after adenosine treatment. Arrows at the top indicate the receptors that most likely mediate the response to MTA and adenosine, with arrow thickness indicating the degree to which these receptors may be involved. However, both MTA and adenosine will each likely interact with multiple types of adenosine receptors to varying degrees and we did not pharmacologically test the impact of adenosine on the A_1A_ and A_3A_ receptors in this model. We saw that A_2B_ receptor antagonism at the time of MTA treatment fully blocked upregulation of *Timp1, Vegfa*, and *Arg1* but did not completely inhibit *IL10* upregulation (Fig. 4b), suggesting MTA signaling is likely being mediated by more than one receptor/pathway, possibly A_3A_, as this receptor has several downstream mediators in common with A_2B_. Similarly, A_2A_ receptor antagonism did not block *IL10* upregulation in response to adenosine treatment, suggesting involvement of another receptor in activating downstream C/EBP signaling. However, we did see upregulation of *IL6* expression following A_2A_ receptor antagonism at the time of adenosine treatment, indicating that the A_2A_ receptor does play a role in the signaling balance mediated by adenosine in this model. Furthermore, the different results from CREB and STAT3 inhibition on IL6 expression at the time of Adenosine or MTA treatment strongly indicate different signaling pathways mediating the response to these metabolites. Gray arrows indicate putative pathways based on previous literature (75-77).

## DISCUSSION

Understanding and targeting the immuno-suppressive mechanisms of GBM is a critical step toward improving treatment for this lethal cancer type (7). In this study, we provide multiple lines of evidence to link MTAP loss, one of the most common genetic/epigenetic events in GBM, to changes in the tumor immune microenvironment. We demonstrate using *in vitro* tumor models that *MTAP* deleted cells have a more anti-inflammatory gene expression signature, and that this immunosuppressive state is recapitulated in the gene expression and immune infiltrate data from GBM patient samples. Loss of MTAP is expected to have profound impact on tumor cells, including conferring susceptibility to PRMT5 inhibition (22,23) and reshaping the methylome landscape and identity of GBM cells (Hansen et al, submitted). Thus, it is not surprising that *MTAP* status also influences the expression of immune regulatory genes within this cell population. The type of differential gene expression analysis we performed on cultured tumor cells, without comprehensively considering non cell-autonomous effects on tumor immune infiltrates, is an obvious limitation. Nevertheless, findings from these *in vitro* models were supported by the results from GBM samples, including differential expression of immune regulatory genes and differential representation of immune cell types.

We discovered through our analysis of GBM samples that one of the cell types profoundly affected by *MTAP* status was the macrophage, with MTAP loss pushing macrophages toward alternative, immune-suppressive M2 polarization. Utilizing *in vitro* models we verified that the process of M2 polarization in this context is influenced by MTA signaling through the adenosine A_2B_ receptor and is dependent on the activity of STAT3, ultimately resulting in upregulation of M2 marker genes *Arg1, IL10*, and *Vegfa*. This represents a unique mechanism of alternative activation and a potentially significant contribution to the immunosuppressive tumor microenvironment in GBM, which is known for an abundance of immunosuppressive tumor-associated macrophages.

It has previously been demonstrated that MTA can directly suppress the proliferation and function of T lymphocytes (24,67). Our findings complement these studies by showing that MTA, which accumulates in MTAP-deficient tumor cells, can also influence innate immune cells. Further research involving *in vivo*, using orthotopic GBM models will be necessary to illuminate the functional interplay between MTAP loss and the function of innate immune cells in GBM, and how they collectively influence the adaptive immune characteristics within the tumor. One question raised by this study is what might be the long term impact of the uptake/accumulation of MTA (i.e., via nucleoside transporters) on the epigenomes and epiproteomes of immune cells, an aspect awaiting further investigation, though previous studies indicate this might play an important role (68-70). Furthermore, we note that the experimental results represent a limited view of the complex time scale of the signaling mechanisms and responses involved. Nevertheless, our findings are revealing, as they unambiguously demonstrate the rapid effect of adenosine receptor A_2B_ signaling initiated by MTA, in a manner distinct from adenosine, on the identity and functionality of macrophages, the most abundant immune cell type in GBM tumors (43).

Importantly, our finding that the macrophage response to MTA is distinct from the response to adenosine suggests that MTAP-deficient GBMs with aberrant MTA accumulation likely have an impact on the immune environment in a manner that is qualitatively and/or quantitatively different from tumors which simply accumulate adenosine as a mechanism of immune evasion (71,72). Using *in vitro* models we found that MTA more potently stimulated expression of M2 macrophage marker genes *Arg1, Vegfa*, and *Timp1* and that MTA and adenosine are both capable of upregulating *IL10* expression through divergent pathways which eventually converge on the transcription factors C/EBP and STAT3, as inhibition of each was able to block upregulation of *IL10* in response to either MTA or adenosine. IL-10 and IL-6 both engage receptors on the cell surface which activate the JAK-STAT signaling pathway. STAT3 is known to positively regulate *Arg1, Vegfa*, and *Timp1*, and we showed that inhibition of STAT3 completely eradicated the upregulation of these genes in response to MTA, confirming a dominant/ obligatory role of STAT3 as a mediator of MTA-stimulated M2 polarization. Additionally, the exclusive upregulation of *IL10* expression by MTA, as opposed to combined *IL6* and *IL10* upregulation initiated by adenosine, is a potential explanation for why MTA results in such an anti-inflammatory gene expression profile, in accordance with the reported opposing forces of IL-6 and IL-10 on STAT3 (64,65) and the potent immunosuppressive effects of IL-10 (66).

The reasons for the different effects of MTA and adenosine on the activation of macrophage marker genes remains unclear. One possible explanation is that these two metabolites engage the various adenosine receptors involved to a different extent. Adenosine is known to have the highest affinity for the A_1_ and A_3_ receptor subtypes, followed by A_2A_ receptors, and has much lower affinity for the A_2B_ receptor by about 50-fold (73). Based on the clear impact of A_2B_ receptor antagonism in mitigating the effects of MTA (but not the effects of adenosine) in our model, one possibility is that MTA either more potently or more exclusively activates A_2B_ than does adenosine (**Fig. 6**), though it is likely that each metabolite exerts some degree of influence on all four adenosine receptor subtypes with varying affinities. In theory, and depending on cell type, the response to MTA or adenosine will include a summarization of purinergic signaling through a combination of these receptors. Activation of G-protein coupled adenosine A_2A_ and A_2B_ receptors invariably results in increased cAMP levels, but each receptor can couple with more than one type of G protein (74) resulting in distinct combinations of downstream signaling pathways (75,76). It is these types of subtle differences in adenosine receptor function which are likely responsible for the different outcomes following MTA or adenosine treatment in this model. Specifically, evidence from our macrophage model showing robust stimulation of *IL6* expression by adenosine that is unaffected by inhibition of A_2B_R and is augmented by inhibition of A_2A_R is indicative of involvement of the A_2A_ receptor and also the A_1_ receptor as likely contributors to the adenosine response in these cells. Though we did not specifically test the involvement of A_1_ or A_3_ receptors in our model, the A_1_ receptor has been shown in HEK293 cells to activate the transcription factor NF-kB (77), a well-documented regulator of *IL6* expression (78-81). We found that inhibition of another transcription factor, C/EBP, which is activated downstream of the A_2A_ receptor, phenocopied A_2A_R inhibition in further upregulating *IL6* expression after adenosine treatment. NF-kB and C/EBP are known to physically associate with one another, resulting in inhibition of promoters with NF-kB enhancer motifs (82). Thus, when considering the potential combinations of receptors involved in generating this response to adenosine, the A_1A_ receptor could lead to activation of NF-kB, and the A_2A_/A_2B_ receptors could lead to activation of C/EBP. Inhibition of A_2A_ receptors or C/EBP would then throw off the balance of these two signaling pathways to favor IL-6 upregulation through NF-kB. We found that MTA, on the other hand, has only a minimal effect on *IL6* expression, which was increased only when combined with a C/EBP inhibitor, and not with a CREB inhibitor or A_2A_ antagonist, supporting the idea that MTA signaling is predominantly mediated through the A_2B_ receptor, which was required for the upregulation of the other M2 polarization gene, and which has been reported as capable of directly inhibiting NF-kB activation (83).

Overall, the complicated roles of adenosine in regulating inflammation, as well as its tumor-promoting functions, are beginning to be more widely recognized (84). Adenosine receptors and the adenosine-generating enzymes, CD39 and CD73, are being investigated as therapeutic targets, either through stimulating or countering the adenosine receptor signaling pathways in order to control pathogenic inflammation or treat advanced cancers (85-87). In GBM, adenosine (and upregulation of CD39 and CD73) has been shown to contribute to immunosuppression (88). Additionally, A_2B_ receptor signaling has been shown to promote tumor metastasis and stimulate tumor angiogenesis (89,90). Our finding that the effect of MTA on macrophage functionality is mediated by the adenosine A_2B_ receptor and is amenable to pharmacological intervention adds a new line of evidence to support the rationale of targeting A_2B_ receptor signaling, in particular in MTAP-deficient GBMs. Development of MTAP-deficient, immune-competent GBM models will be necessary for further investigating the effectiveness of this strategy *in vivo*. We speculate in the case of *MTAP*-null GBM, given the abundant presence of tumor-associated macrophages and the characteristic immunosuppressive microenvironment, that the effect of A_2B_ antagonists will be a particularly relevant area for investigation, with the potential to significantly impact tumor growth and augment immunotherapeutic interventions in *MTAP*-deleted GBM.

## MATERIALS AND METHODS

### Cell lines and cell culture

RAW 264.7 and THP-1 macrophage cell lines were obtained from the Duke Cell culture Facility. BV-2 cells were a generous gift from Dr. Tso-Pang Yao. RAW 264.7 and BV-2 cells were maintained in DMEM with 4.5g/L L-glucose (Sigma Cat #D6429) supplemented with 10% heat-inactivated FBS and anti-anti (antibiotic/antimycotic). THP-1 cells were cultured in RPMI 1640 with 10% heat-inactivated FBS and anti-anti. CT-2A cells were a generous donation from Dr. Darrell Bigner. The cells were maintained in DMEM/F12 (Gibco Cat #11330-032) supplemented with B-27 (Gibco Cat #17504-044), EGF (Stemcell), and FGF (Stemcell) and grown in suspension. Primary tissue cultures were derived with consent from patient tumor samples obtained by the Duke Brain Tumor Center. These patient-derived cultures were maintained in human neural stem cell (NSC) media (STEMCELL, cat# 05751), supplemented with EGF, FGF, and Heparin and plated onto laminin coated plates. All experiments were performed within the first 20 passages.

### Plasmid construction and generation of derivative cell populations

The CRISPR system was used for knockout of *Mtap* in CT-2A. Two double-stranded oligonucleotides that encode sgRNA targeting exon 1 and exon 3 of *Mtap* were cloned into the px552-pEASY plasmid. CT-2A cells were transfected simultaneously with sgRNA plasmid and the px458 plasmid containing cas9 and GFP. For transient plasmid transfection, plasmids (2 plasmids at 1:1 ratio for achieving the desired gene deletion/mutations) and Transfex (ATCC, cat# ACS-4005) were mixed and used for cell transfection according to manufacturer’s instructions. Three to four days after the transfection, green fluorescent protein–positive (GFP+) cells were sorted via fluorescence-activated cell sorting (BD FACSVantage SE cell sorter, Duke Cancer Institute) to obtain the GFP+ population. Sorted cells were plated at single-cell densities and allowed to expand for 21 days, at which point DNA was prepped from each colony to screen for a deletion in *MTAP* (exon 1-exon 3) using PCR amplification across the deleted region. sgRNA and primer sequences are shown in Supporting Material Table 1.

### In vitro macrophage polarization

Raw 264.7 cells were plated in a 12 well dish and stimulated with methylthioadenosine (MTA) (Cayman Cat #15593) or adenosine (Sigma Cat #A9251) for 12 hours, at which point they were collected for analysis. Cytokines IL4 (Peprotech Cat #200-04) and IL-13 (Peprotech Cat #200-13) were added within one hour of MTA/adenosine administration at a dose of 5ng/mL. THP-1 stimulation with MTA was identical to RAW 264.7 cells except that the cells were first differentiated using PMA (Cayman Cat #10008014) for 24 hours, then cultured in fresh media for 24 hours prior to stimulation. To test the impact of A_2B_ and A_2A_ receptors on MTA-mediated macrophage polarization, A_2A_R inhibitor Istradefylline (Selleckchem Cat #S2790) or A_2B_R inhibitor PSB0778 (Tocris Cat #3199) were added simultaneously with MTA or adenosine, except in the indicated experiments where delayed administration of the A_2B_R inhibitor was tested. To further explore downstream signaling pathways STAT3 inhibitor Stattic (Tocris Cat # 2798), CREB inhibitor 666-15 (Tocris, Cat #5661) and C/EBP inhibitor betulinic acid (Tocris Cat #3906) were added at the time of MTA/adenosine administration. To test the effect of physiological accumulations of MTA from tumor cells, spent media from CT-2A parental or MTAP knockout cells was collected and added to the macrophage media in a 2:1 ratio and cells were plated for 12 hours prior to measuring response.

### Preparation of RNA and RT-qPCR

Total RNA was extracted using quick-RNA mini prep kit (Zymo Research, cat# 11-328) following the manufacturer’s protocols. Concentration of RNA was determined by Nanodrop Lite Spectrophotometer (Thermo Scientific). For gene expression analysis, reverse transcription was performed to convert total RNA into complementary DNA (cDNA) using the RNA to cDNA EcoDry Premix (Clontech, cat #639547). Subsequently, real-time qPCR was performed following the aforementioned qPCR procedure. Each reaction included a cDNA template equivalent of 10 ng of total RNA. The glyceraldehyde 3-phosphate dehydrogenase (*GAPDH*) and beta-Actin genes were used as internal expression controls for RT-qPCR with reliable results. When beta Actin was used as the control amplicon the following program was followed: 95°C, 3 minutes; 41 cycles of 95°C 10 seconds and 68°C 20 seconds, then a standard dissociation curve from 65°C to 95°C of 5 seconds/ 5 degree increment. When Gapdh was utilized as the internal control the following program was used: 95°C, 3 minutes; 40 cycles of 95°C 10 seconds, 60°C 20 seconds, and 72°C 1 second, then a standard dissociation curve from 65°C to 95°C of 5 seconds/ 5 degree increment.

### Oligos and primers

All oligos and primers used for the study were synthesized by Eton Bio and are listed in **Supporting data Table 1**. Quantitative PCR was performed using KAPA SYBR Fast 2x Universal master mix (KK4602) according to the manufacturer’s protocols on a BIO-RAD CFX96 Real-Time System.

### Gene expression microarray

The RNA was extracted as described above. Samples were analyzed using the Affymetrix Human Genome U133 Plus 2.0 array according to the manufacturer’s protocols by the Duke Sequencing and Genomic Technologies Shared Resource. Data was analyzed using the Affymetrix Expression Console and Affymetrix Transcriptome Analysis Console v3.0 software.

### Pathway analysis

Pathway analysis was done using the DAVID 6.8 platform at david.ncifcrf.gov (91,92). For pathway analysis of gene expression differences between *in vitro* samples and between patient cohorts, all genes found to be “significant” by the parameters defined in the manuscript were included in the analysis, with a maximum of 3000 genes allowed by DAVID (if more than 3,000 probes showed significantly different expression levels between the two populations, those with the highest difference were included).

### Gene Set Enrichment Analysis

Gene set enrichment analysis was performed using the GSEA 3.0 software as previously described. Custom gene sets were generated containing HLA genes and inflammatory signaling molecules. Gene sets are listed in supporting information table 2.

### Analysis of TCGA data

All TCGA data was downloaded from the online portal https://tcga-data.nci.nih.gov/docs/publications/tcga/ and through cbioportal.org (33,34). The most recently published 2013 GBM data set was used for all analyses. For each analysis, the maximum number of complete cases available (confirmed *IDH1/2* wildtype) were used unless otherwise stated, as IDH mutations are known to independently influence epigenetics and cellular differentiation. Analyses performed include gene expression (385 samples). We utilized *MTAP* expression levels to categorize patients rather than gene copy number because *MTAP* is known to be silenced epigenetically in a variety of cancer types (93-96), and our analysis (Hansen et al, submitted) of DNA methylation and gene expression data in patients suggests it can also be epigenetically silenced in GBM. To perform this analysis, samples were equally divided into quartiles based on MTAP expression and a *t* Test was used to compare the “low” and “high” groups to find differences in gene expression between these two groups.

### Statistical analysis

Statistical tests (student’s t test, ANOVA) were performed using Graphpad Prism. All experiments were repeated to ensure reproducibility of results. Unless otherwise indicated, pooled data from multiple experiments was used for each figure. A *P* value cutoff of 0.05 was used to determine significance in all cases except where corrections were applied for larger data sets (ie. Bonferonni).

## Acknowledgements

We thank Ping Fan of the Duke Cancer Institute’s Pharmacokinetic/Pharmacodynamic core laboratory for the LC-MS/MS metabolite analysis, and Heather Hemric, Laura-Leigh Rowlette, and Holly Dressman of the Duke Sequencing and Genomic Technologies Shared Resource for the Affymetrix array processing service.

## Author Contributions

**Conception and design:** L.J. Hansen, Y. He

**Methodology development:** L.J. Hansen, R. Yang, Y. He

**Data acquisition:** L.J. Hansen, R. Yang, K. Woroniecka

**Data analysis and interpretation:** L.J. Hansen, K. Woroniecka, Y. He

**Writing, reviewing, and revision of the manuscript:** L.J. Hansen, R. Yang, K. Woroniecka, H. Yan, Y. He

**Administrative, technical, or material support:** H. Yan, Y. He

**Study supervision:** Y. He

## FOOTNOTES

This work was supported by a National Cancer Institute National Research Service Award (F30CA206336; L.J.H), a National Comprehensive Cancer Network Young Investigator Award (Y.H.), and the National Institute Of Neurological Disorders And Stroke of the National Institutes of Health, Award Number R01NS101074 (Y.H). This work was also supported by funding from the National Institutes of Health Duke SPORE in Brain Cancer (P50 CA190991), as well as the Circle of Service Foundation (Y.H) and the Southeastern Brain Tumor Foundation (Y.H).

## The abbreviations

MTA: methylthioadenosine
MTAP: methylthioadenosine phosphorylase
GBM: glioblastoma
RT-qPCR: reverse transcription quantitative polymerase chain reaction
B. Acid: Betulinic Acid
LC-MS/MS: Liquid chromatography tandem mass spectrometry

